# Identifying and validating the presence of Guanine-Quadruplexes (G4) within the blood fluke parasite *Schistosoma mansoni*

**DOI:** 10.1101/2020.09.09.289975

**Authors:** Holly M Craven, Riccardo Bonsignore, Vasilis Lenis, Nicolo Santi, Daniel Berrar, Martin Swain, Helen Whiteland, Angela Casini, Karl F Hoffmann

## Abstract

**Background:** Schistosomiasis is a neglected tropical disease that currently affects over 250 million individuals worldwide. In the absence of an immunoprophylactic vaccine and the recognition that mono-chemotherapeutic control of schistosomiasis by praziquantel has limitations, new strategies for managing disease burden are urgently needed. A better understanding of schistosome biology could identify previously undocumented areas suitable for the development of novel interventions.

**Methodology/Principal findings:** Here, for the first time, we detail the presence of G-quadruplexes (G4) and putative quadruplex forming sequences (PQS) within the *Schistosoma mansoni* genome. We find that G4 are present in both intragenic and intergenic regions of the seven autosomes as well as the sex-defining allosome pair. Amongst intragenic regions, G4 are particularly enriched in 3’ UTR regions. Gene Ontology (GO) term analysis evidenced significant G4 enrichment in the *wnt* signalling pathway (*p*<0.05) and PQS oligonucleotides synthetically derived from *wnt*-related genes resolve into parallel and hybrid G4 motifs as elucidated by circular dichroism (CD) spectroscopy. Finally, utilising a single chain anti-G4 antibody called BG4, we confirm the *in situ* presence of G4 within both adult female and male worm nuclei.

**Conclusion/Significance:** These results collectively suggest that G4-targeted compounds could be tested as novel anthelmintic agents and highlights the possibility that G4-stabilizing molecules could be progressed as candidates for the treatment of schistosomiasis.

**Author Summary:** *Schistosoma mansoni* causes schistosomiasis, a parasitic disease that affects millions of people living in resource-deprived areas of developing countries. No vaccine exists and the current drug treatment has limitations, notably inefficacy against the larval stages of the parasite. New drugs are, therefore, needed to sustainably control schistosomiasis. A further understanding of parasite biology will uncover new targets and lead to the development of novel therapies. Here, we identify the presence of G-Quadruplexes (G4s) in *S. mansoni*. G4s are four-stranded DNA structures that can affect gene function and, to date, have not been previously found in any parasitic helminth. Computational analysis predicted potential G4 folding sequences within the *S. mansoni* genome, several of which were confirmed to fold by circular dichroism spectroscopy. Analysis of G4-containing protein coding genes found an enrichment within the *wnt* signalling pathway, a developmental pathway crucial for axial development in the parasite. Additionally, G4s could be detected within adult worms using a fluorescent antibody that selectively recognises quadruplex structures in nucleic acids. This research describes the presence of a previously unknown structure within the parasite, which could present a new target for developing novel treatments.

## Introduction

*Schistosoma mansoni*, a digenean platyhelminth responsible for the neglected tropical disease (NTD) schistosomiasis, maintains its complex lifecycle through definitive human and intermediate snail (*Biomphalaria sp*.) hosts. Exhibiting dioecy as adults, mature *S. mansoni* pairs establish infection within the human mesenteric vessels draining the intestine. While adults are largely non-immunogenic, copulation between schistosome pairs leads to the production of an estimated 300 immunogenic eggs per day. Approximately half of these eggs migrate through the mesenteric veins and reach the intestinal lumen where they are released with faeces, a requirement for lifecycle transmission. Endemic in sub-tropical regions both in the new and old world, schistosomiasis contributes to socioeconomic developmental arrest and is a public health burden in many developing countries. It is estimated that some 250 million individuals worldwide suffer from schistosomiasis per annum, with the disease accounting for the loss of 2.5 million DALYs (disability adjusted life years) in 2016 [1].

The *S. mansoni* genome, first published in 2009 [2], updated in 2012 [3] and under continual refinement in WomBase-Parasite (current assembly version 7) has facilitated the advancement of many basic investigations of schistosome biology [4–6]. However, much of what powers survival and developmental success within the parasite remains unknown. It is, therefore, vital that research identifying vulnerabilities in the schistosome genome continues to facilitate development of new schistosomiasis control strategies in light of existing limitations associated with praziquantel mono-chemotherapy [7,8].

Guanine quadruplexes (G4s) have recently been implicated as targets for the control of both infectious diseases (caused by *Plasmodium, Leishmania* and *Trypanosoma* species) and noncommunicable diseases (notably in solid cancers) [9–14]. Found to be evolutionarily conserved across phyla (eukaryotes, prokaryotes, viruses), these four stranded DNA (and RNA) structures fold from guanine (G) rich regions in the genome [15–19]. Guanines form planar tetrads mediated through Hoogsteen hydrogen bonds internally stabilised by monovalent cations, usually K^+^ [20]. These tetrads stack and arrange into a tertiary structure known as a quadruplex, joined by intermediate loops formed of any other nucleobase (i.e. cytosine, adenine, guanine, thymine or uracil). While G4s are less stable than duplex DNA and are frequently found in single stranded telomeric overhangs, upstream promoter regions or within genes, their transient formation at these positions can directly affect transcription, translation and replication [21–23]. Approximately 50% of all human genes are predicted to contain a G4 within or near promotor regions [24] as well as in 5’ UTRs [19].

In *P. falciparum*, a species with a particularly AT rich genome, putative quadruplex forming sequences (PQS) are found within VAR genes as well as in telomeric regions [9,13,16]. In *T. brucei*, a main G4 sequence of interest is a highly repeated 29 nucleotide long PQS (referred to as EBR1; Efres Belmonte-Reche-1) which exhibits selective binding to G4 ligand carbohydrate naphthalene diimide derivatives when compared to dsDNA. These ligands were shown to have antiparasitic activity, indicating G4 targeting could be a potential therapeutic avenue [25]. Nevertheless, to our knowledge there have been no in-depth investigations into the presence of G4s within extracellular endoparasites such as those species contained within the Platyhelminthes. As *S. mansoni* possesses a more GC rich genome than *P. falciparum* (∼34% compared to ∼18% respectively) [3,26], this suggests that PQS could be present. Moreover, similar to *P. falciparum, S. mansoni* contains a TTAGGG telomeric tandem repeat, which, in humans, has been demonstrated to fold into G4 and is linked with telomere capping [27]. The discovery of G4s within the schistosome genome could, therefore, identify a new molecular regulator of biological processes and genome architecture. In turn, G4-targetting compounds could modulate these activities and structures, ultimately leading to the development of novel strategies for controlling a major NTD. Here, we present the first description of PQS within *S. mansoni* that will foster further investigations in this promising area.

## Materials and Methods

### Ethics statement

All procedures performed on mice adhered to the United Kingdom Home Office Animals (Scientific Procedures) Act of 1986 (project licenses PPL 40/3700 and P3B8C46FD) as well as the European Union Animals Directive 2010/63/EU and were approved by Aberystwyth University’s (AU) Animal Welfare and Ethical Review Body (AWERB).

### Genome preparation

FASTA files of the *S. mansoni* genome (version 7.1) used for computational analyses of PQS were obtained from the Wellcome Sanger Institute [28]. Scaffolds not assigned to the chromosome assembly were not used. Gene annotations, obtained from the associated general feature format (GFF) extension files, were used to assign PQS to intragenic regions including CDS (coding sequence), five prime UTR (5’ UTR) and three prime UTR (3’ UTR) as defined from the annotation GFF file. Intronic regions were not labelled within the GFF and were calculated by extracting the regions between exons using in house scripts. All other annotation types were excluded.

### Quadparser (QP) analysis of genome

QP [29] was compiled in Linux. Genome FASTA files were run through QP using parameters G_3+_N_1-7_G_3+_N_1-7_G_3+_N_1-7_G_3+_ and C_3+_N_1-7_C_3+_N_1-7_C_3+_N_1-7_C_3+_ where G = guanine, C = cytosine and N = A, T, C or G. This allows G4s present on both the coding and non-coding strand to be detected. Output of PQS were formatted into a browser extensible data (.BED) format file for visualisation with the Integrative Genomics Viewer (IGV) v2.4.2 desktop application [30].

### G4Hunter (G4h) analysis of genome

G4h [31] was executed through a Bourne-again shell (Bash) environment. Genome FASTA were run through G4h using a fixed nucleotide window of 30 and a score threshold of ±1.4. Positive scores indicate G4 on coding strand, and negative scores indicate G4 on the non-coding strand.

### PQS overlap, normalisation and density quantification

The Bedtools package [32] was used to find overlapping PQS in both QP and G4h outputs. These commonly identified loci were then further defined by intragenic occurrences (present in CDS, 5’ UTR, 3’ UTR and intron GFF notations). Bedtools was subsequently used to acquire *S. mansoni* protein coding genes (Smps) containing intragenic PQS. Further Smp data were extracted using the WormBase Parasite Biomart function as well as the scientific literature [2,3,33,34]. PQS density was calculated as the number of PQS per chromosome per Mb of feature/chromosome. A baseline of total PQS density per genome length was calculated to identify over or under enrichment of PQS within intragenic features on different chromosomes. Analysis of enrichment was performed by calculating PQS/Mb densities. The significance of the differences was assessed by χ^2^ test in R [35]. The *p* vales were corrected for multiple testing with the Bonferroni-Holm method [36].

### Gene Ontology (GO) classification

GOAtools analysis scripts [37] were used to quantify functional groupings within the PQS-containing Smps; a *p* value cut-off of 0.05 was selected to identify statistically significant terms. GOAtools analysis was corroborated by using ReVigo under standard settings [38].

### Circular Dichroism (CD) spectroscopy

DNA oligonucleotides (Table 1) for CD analysis were obtained from Eurogentec (Belgium), supplied at 1000 nmol scale, RP-HPLC purified, desalted and reconstituted to 100 µM in biological grade water (VWR, US). Oligos were adjusted to a 6 µM working solution in either 60 mM TrisKCl (10 mM TrisHCl, 50 mM KCl; pH 7.4) or 10 mM TrisHCl (pH 7.4) buffers. Oligos were denatured for 5 min at 95 °C in an oil bath and left to cool to room temperature (RT) prior to analysis. Spectra were recorded on a Chirascan (Applied Photophysics, UK) CD spectrometer measuring between 210-400 nm at 25 °C, step 1 nm. Triplicate reads were performed, and traces averaged and smoothed. A 60 mM TrisKCl (pH 7.4) or a 10 mM TrisHCl (pH 7.4) buffer only background were additionally run, and baseline subtracted from each respective read. For G4-melting assays, the oligonucleotide concentrations were adjusted to 6 µM in 60 mM TrisKCl (pH 7.4) buffer. Spectra were recorded as defined above, with a 5 °C ramp from 25 °C to 95 °C with a 2 min equilibration hold prior to reading.

**Table 1:**
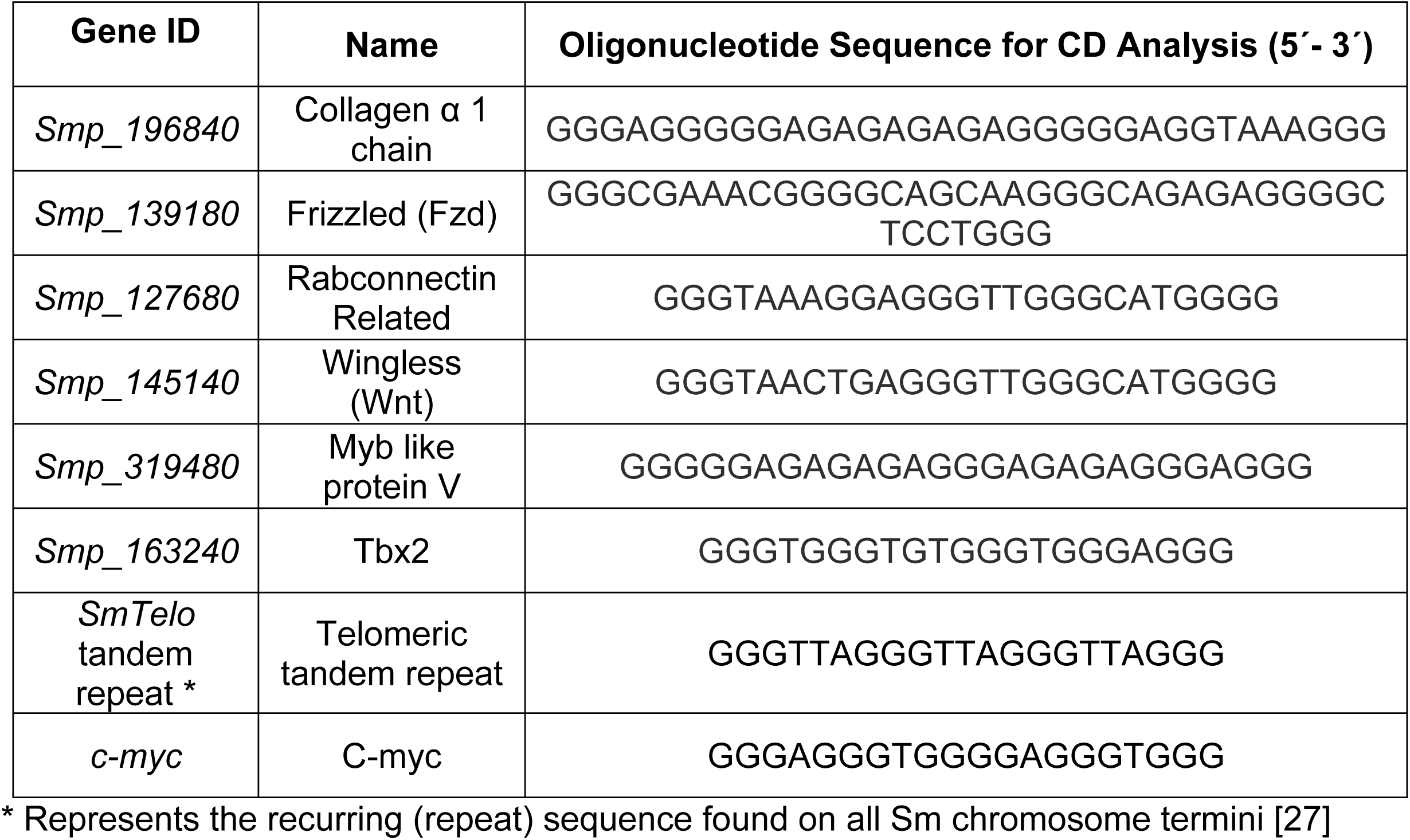
Genomic identifiers and oligonucleotide sequences used in CD spectroscopy.

### *S. mansoni* adult worm cultures

Female HsdOla:TO mice (ENVIGO, UK) were infected with 180 cercariae via percutaneous exposure for 40 min. Adult worms were perfused 7 wk post infection as described previously [39,40]. Parasites were cultured for 24 hrs at a density of three worm pairs per well in a 48 well tissue culture plate (Fischer Scientific, UK) in 1 mL of complete media (DMEM, 1% Penicillin-Streptomycin, 200 mM L-glutamine, 10% FBS, 10% HEPES) at 37 °C in a humidified environment containing 5% CO_2_.

### *In situ* detection of G4s in schistosomes

Cultured 7 wk adult schistosomes were incubated in a 0.25% (w/v) solution of the anaesthetic ethyl 3-aminobenzoate methanesulfonate in complete media for 10 min before being killed in a 0.6 M MgCl_2_ solution. Worms were subsequently fixed overnight in ice cold 3:1 methanol: acetic acid at −20°C prior to 1 hr permeabilisation in PBS containing 0.3% (v/v) Triton X-100 (PBSTx) at RT. Following permeabilisation, schistosomes were blocked for 2 hrs at RT in blocking buffer (5% BSA (w/v) in PBSTx) and then incubated for 72 hrs at 4°C in 1:400 anti- G-Quadruplex antibody BG4 (Ab00174-10.6, Absolute Antibody, Oxford UK) diluted in blocking buffer. Parasites were washed thrice for 15 min in PBSTx and incubated for 24 hrs in 1:2000 F_ab_ specific FITC conjugated goat anti human IgG (F5512, Sigma-Aldrich UK) diluted in blocking buffer. Samples were washed again with PBSTx and mounted on microscope slides in Vectashield anti-fade mounting medium counterstained with 5 μg/mL DAPI. Negative controls were incubated in either 120 U DNase I or 0.1 mg/mL RNase A in PBS for 1 hour at 37°C 5% CO_2_ in a humidified environment post permeabilisation. Slides were imaged on a Leica TCS SP8 Laser Confocal Scanning Microscope at x20 air or x100 oil immersion at 8000 Hz, using DPSS and Argon laser intensities at 20%. Anterior, midsection and posterior images were collated from 100 Z stacks and representative images taken from each group (*n* = 8) are shown. RNase was tested by incubation of 0.1 mg/mL RNase A with 1 µM of either universal mouse RNA (QS0640, ThermoFisher UK) or extracted *S. mansoni* RNA [41] for 1 hour at 37°C. The digested RNA was then electrophoresed on a 1% agarose gel in TAE buffer at 8 V/cm and visualised using a UVP Gel Doc-it.

## Results

The *S. mansoni* genome was searched for the presence of G4 using two different algorithms, quadparser (QP) and G4Hunter (G4h). PQS-containing loci identified by both algorithms were retrieved and used to compile a high stringency list of overlapping genomic positions (*n* = 406) (Fig. 1). While high stringency searches will result in fewer returned sequences, they have the greatest likelihood of folding canonical intramolecular G4 *in vivo*. Here, high stringency selected PQS were found unequally distributed across both intragenic (*n* = 122) and intergenic (*n* = 284) regions.

**Figure 1.**
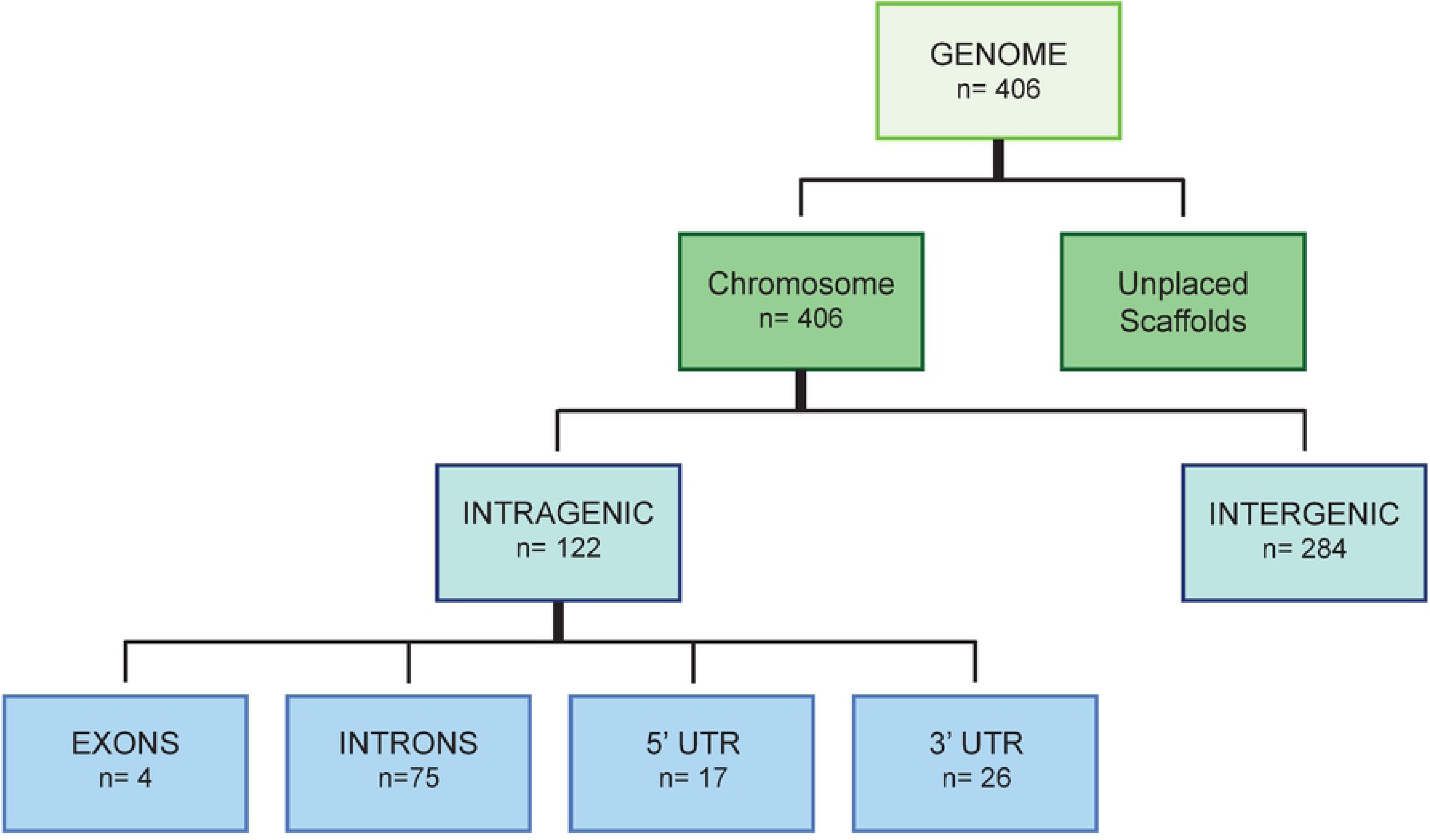
Flowchart for identifying PQS found in the *S. mansoni* genome. Quadparser and G4Hunter were used to independently identify PGS within the *S. mansoni* genome (v7.1). The Bedtools intersect function was subsequently used to find overlapping PQS detected by both methods. Across the assembled chromosome scaffolds (genome), this led to the discovery of 406 PQS. Unplaced scaffolds were not included in the analysis. Intragenic PQS analysis, determined using the genome annotation file, found 122 of these to be within either Exons (4), Introns (75), 5’ UTRs (17) or 3’ UTRs (26). Those PQS deemed intergenic are found within chromosomes but not within intragenic regions.

Global distribution of these selected 406 high stringency PQS (both intragenic and intergenic) across the *S. mansoni* genome was first assessed (Fig. 2). Initial observations indicated that PQS were found within all seven autosomes as well as the sex-defining allosome ZW pair, with numerous PQS appearing towards the ends of chromosomes (Fig. 2A). Furthermore, PQS were identified on both template and non-template strands as well as at chromosomal termini in the form of TTAGGG tandem telomeric repeats on all chromosomes (Fig. 2B, S1 Table) [27]. The largest total number of PQS was found on chromosome 5 (n=100) and the smallest number was found on chromosome 7 (*n* = 19) (Fig. 2C and Table 2).

**Table 2:** PQS numbers and locations within each *S. mansoni* chromosome

| Chromosome | Total Number of PQS and Location Within Each Chromosome |  |  |  |  |  |
| --- | --- | --- | --- | --- | --- | --- |
|  | All PQS | All intragenic PQS | Intragenic PQS |  |  |  |
|  |  |  | Exon | 5' UTR | 3' UTR | Intron |
| 1 | 48 | 10 | 2 | 3 | 2 | 12 |
| 2 | 59 | 9 | 0 | 4 | 2 | 14 |
| 3 | 54 | 17 | 1 | 4 | 3 | 7 |
| 4 | 28 | 8 | 1 | 2 | 1 | 7 |
| 5 | 100 | 7 | 0 | 0 | 6 | 6 |
| 6 | 46 | 13 | 0 | 1 | 4 | 10 |
| 7 | 19 | 9 | 0 | 0 | 3 | 4 |
| ZW | 52 | 22 | 0 | 3 | 5 | 15 |
| **Total** | 406 | 122 | 4 | 17 | 26 | 75 |

**Figure 2.**
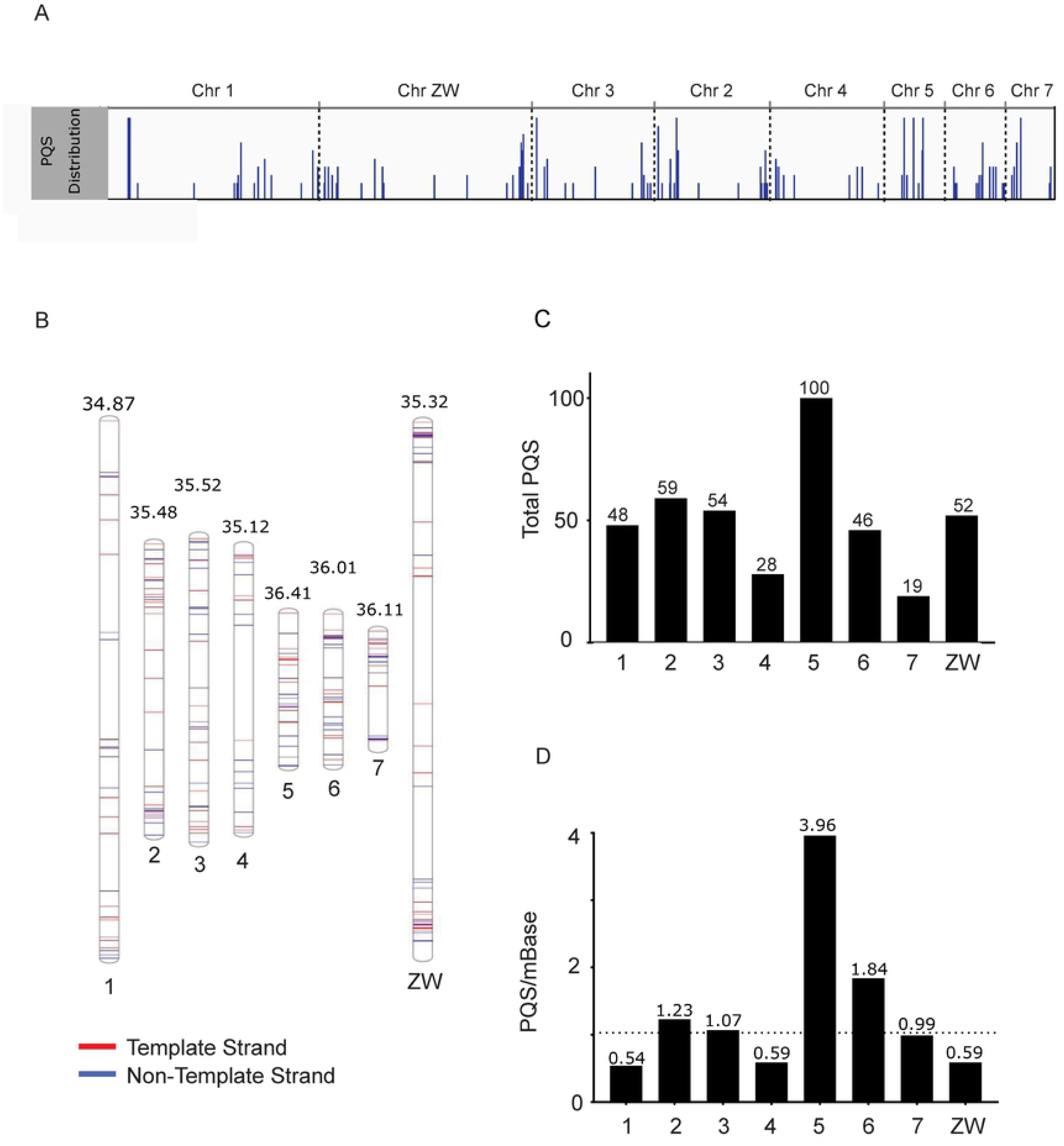
The distribution of PQS across the *S. mansoni* chromosomal assembly. Both intragenic and intergenic PQS (*n* = 406) were mapped to chromosomes (*S. mansoni* genome, v7.1). A) IGV snapshot of the chromosomal assembly, where PQS appearing in close proximity are visualised as longer bars. Chromosomes are ordered according to size (largest to smallest). B) A phenogram mapping the position of PQS along the length of the chromosome (autosomes ordered sequentially). PQS appear on both forward and reverse strands. Numbers above each chromosome represent % GC content. C) Total number of PQS identified per chromosome (sequentially ordered). Chromosome 5 has the greatest number of PQS (*n* = 100) whilst the fewest were detected in chromosome 7 (*n* = 19). D) PQS density per Mb of chromosome. Average PQS/Mb found in the complete chromosomal assembly is indicated by a dashed line and denotes a baseline. Enrichment of PQS within any chromosome is found above this dotted line.

To test if PQS are randomly distributed or are enriched in particular chromosomes, PQS per Mb were calculated for each chromosome and compared to a baseline (average of 1.03 PQS per Mb found in the complete chromosomal assembly, Fig. 2D). Data points above the baseline can be considered enriched in PQS (Fig. 2D and Table 3). Accordingly, four *S. mansoni* chromosomes (chromosomes 2, 3, 5 and 6) are enriched for PQS when compared to the baseline frequency detected across the whole genome, with chromosome 5 particularly enriched. Chromosomes 1, 4, 7 and ZW were considered unenriched for PQS by this analysis. After adjusting for chromosome size, we used χ^2^ test to assess the PQS distribution across normalised chromosomes and observed a significant difference (*p* < 2.2×10^−16^). We then carried out all pair-wise comparisons using the Bonferroni-Holm method to adjust for multiple testing. We found that chromosome 5 had a greater density of total PQS, compared to any other chromosome (3.96 PQS/Mb). Given chromosome 5 is a relatively short chromosome (25 Mb), this indicated that the presence of PQS is not solely driven by chromosome length and enrichment may be driven by specific chromosome features, such as intragenic motifs. The effect of chromosome GC content on total and intragenic PQS content was assessed by regression analysis and found to have no association (R^2^ = 0.14 for total PQS and R^2^ = 0.29 for Intragenic PQS).

**Table 3:** PQS density characteristics (PQS/Mb) within each *S. mansoni* chromosome

| Chromosome | PQS per Mb of feature length |  |  |  |  |  |  |
| --- | --- | --- | --- | --- | --- | --- | --- |
|  | Chr Length (Mb) | All PQS | All Intragenic PQS | Intragenic |  |  |  |
|  |  |  |  | Exon | 5' UTR | 3' UTR | Intron |
| 1 | 88.9 | 0.55 | 0.37 | 0.48 | 0.44 | 0.54 | 0.33 |
| 2 | 48.1 | 1.23 | 0.76 | 0 | 1.22 | 1.23 | 0.73 |
| 3 | 50.5 | 1.07 | 0.58 | 0.46 | 1.22 | 1.64 | 0.38 |
| 4 | 47.3 | 0.59 | 0.43 | 0.48 | 0.56 | 0.62 | 0.38 |
| 5 | 25.3 | 3.96 | 1.01 | 0 | 0 | 7.69 | 0.68 |
| 6 | 25.0 | 1.84 | 1.08 | 0 | 0.54 | 4.49 | 0.99 |
| 7 | 19.3 | 0.99 | 0.6 | 0 | 0 | 4.23 | 0.45 |
| ZW | 88.4 | 0.59 | 0.44 | 0 | 0.44 | 1.27 | 0.41 |
| Baseline | - | 1.03 | 0.56 | 0.23 | 0.60 | 1.73 | 0.48 |

We reasoned that a more thorough interrogation of PQS found only within intragenic regions would generate further clues for interpreting PQS function in the *S. mansoni* genome. Therefore, PQS were next classified according to their location within intragenic features (defined here as 5’- UTR, 3’- UTR, Exon and Intron) of annotated Smps (Fig. 3, Table 2 and Fig. 1). Analysis of these features concluded that 30% of the 406 PQS found within the *S. mansoni* genome were located within genes (n = 122) with most found in the introns (n = 75) and fewest found in exons (n = 4) (Fig. 3A and Fig. 1). Furthermore, this analysis indicated that distribution of PQS across intragenic features is not equal: chromosome ZW had the highest number of intragenic PQS (n=23) and chromosome 7 contained the fewest PQS (n = 7). We observed a significant difference in the density distribution of PQS across the four intragenic regions (*p* = 3.73×10^−9^, χ^2^ test), with the highest density of 1.76 PQS per Mb found on 3’ UTR. By contrast, exons had the lowest density, with 0.23 PQS per Mb (Fig. 3B and Table 3). A post-hoc pair-wise comparison with Bonferroni-Holm correction for multiple testing revealed that, relative to the size of the intragenic regions, there were significantly more PQS found on 3’ - UTR compared to 5’- UTR (p = 0.0029), exon (p = 9.3×10^−5^) and intron (p = 2.6×10^−8^) features.

**Figure 3.**
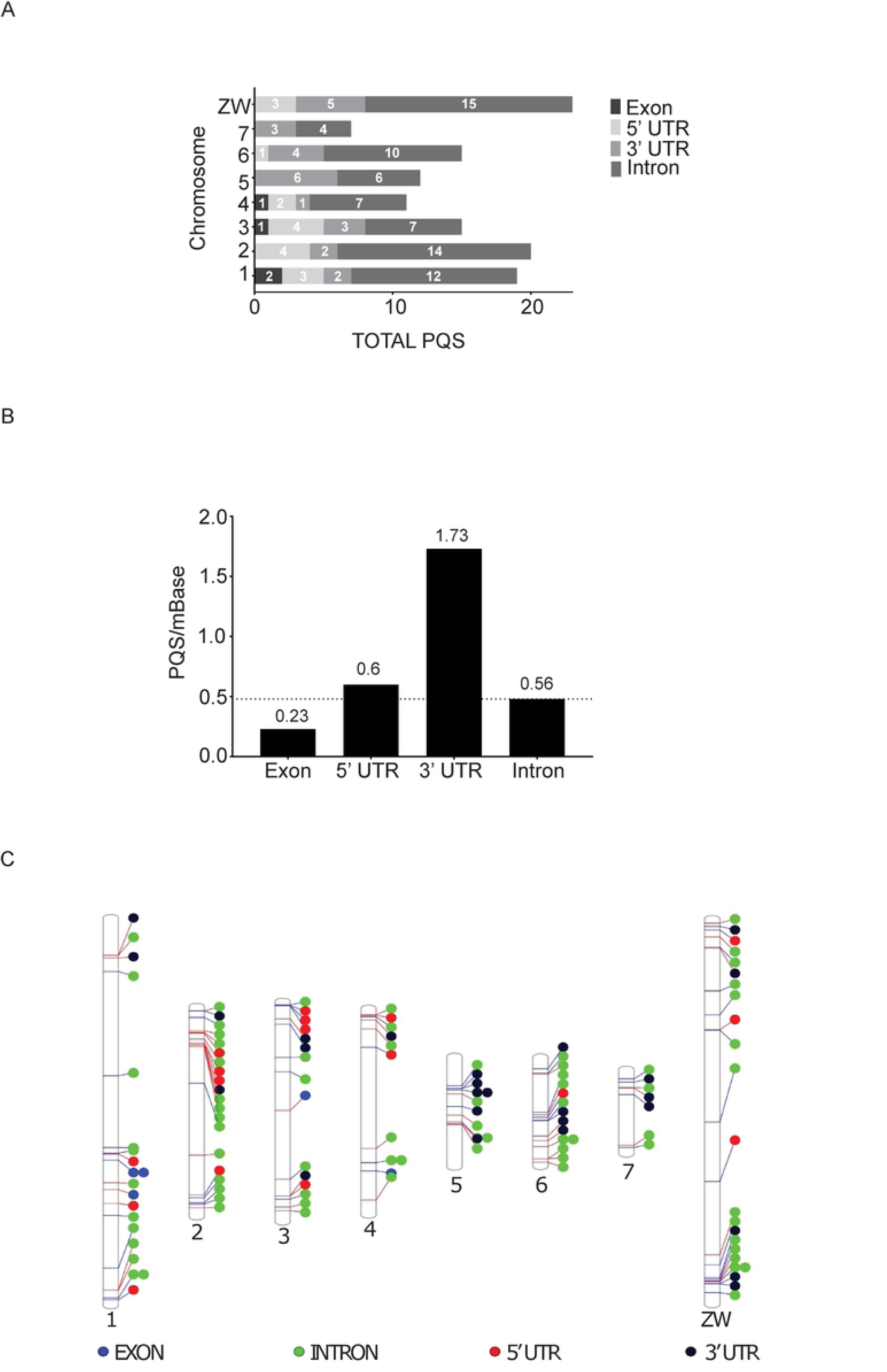
Intragenic PQS distribution within the *S. mansoni* genome. A) The distribution of intragenic PQS per individual chromosomes. For each chromosome, intragenic PQS were categorised into those found within introns, exons, 5’- and 3’- UTRs. The largest number of PQS were found on chromosome ZW and the fewest on chromosome 7, with most PQS aligned to intronic features and fewest PQS found within exons. B) Intragenic PQS per Mb were calculated for each chromosomal feature and compared to the average intragenic PQS per Mb for all features (dotted horizontal line = baseline). Both 5’- and 3’- UTR categories had greater PQS per Mb than the calculated baseline (0.56). Multiple comparisons found 3’- UTRs were enriched with more PQS/mb compared to other features. C) Phenogram detailing localisation of intragenic PQS in each chromosome. Coloured circles denote PQS location according to intragenic feature.

PQS distribution across features was further evaluated at the chromosome level to identify where each of the intragenic PQS were located (Fig. 3C and Table 3). Here, intragenic PQS were found broadly distributed across each of the chromosomes. GO term enrichment analysis was subsequently performed on all unique Smps (i.e. some Smps contained more than one PQS) containing PQS (n=106) using GOAtools [37] and ReVigo [38]; the most significant enrichment in GO terms 0016055 (*wnt* signalling pathway) and 1905114 (cell signalling) (*p* < 0.00002) were found (S1 Fig.).

Following genome analysis, several PQS-containing Smps (Table 1) were selected for circular dichroism (CD) spectroscopy to investigate their folding *in vitro* [42,43]. An oligonucleotide representing *smTelo* was further selected as a control for G4 antiparallel/hybrid folding, as it contains an identical sequence to the well-studied *Homo sapiens* tandem telomeric motif TTAGGG (herein referred to as *hTelo*) located at chromosome termini found to adopt this particular G4 structure [44]. Finally, an oligonucleotide representing *H. sapiens c-myc* was utilised as a control for the formation of G4 parallel folding [45].

In the presence of K^+^, five (*smp_139180, smp_145140 smp_163240, smp_319480, smp_127680*) of the six PQS-containing oligonucleotides selected showed a CD spectra compatible with G4-folding (Fig. 4). When compared with the *c-myc* G4 sequence, the spectra of *smp_139180 (frizzled), smp_145140 (wnt)* and *smp_163240 (tbx2)* all produced traces with the same positive peaks (between 262 nm – 263 nm) and negative troughs (between 241 nm – 242 nm) characteristic of a parallel G4 structure (Fig. 4A). PQS found in *smp_319480* (*myb like protein*) and *smp_127680* (*rab connectin*) displayed a positive peak at 263 nm, a negative peak at 238 nm and a shoulder at 293 nm, mirroring the spectra observed for *smTelo* (a negative peak at 238 nm, shoulder peak at 263 nm and positive peak at 293 nm) suggesting their formation of an antiparallel hybrid quadruplex conformation (Fig. 4B) [42,46]. In contrast, the CD spectrum of the PQS identified in *smp_196840 (collagen α 1 chain*) could not be associated with any known G4-CD profiles, leading us to exclude its folding into any G4 structure in the applied experimental conditions (Fig. 4C).

**Figure 4.**
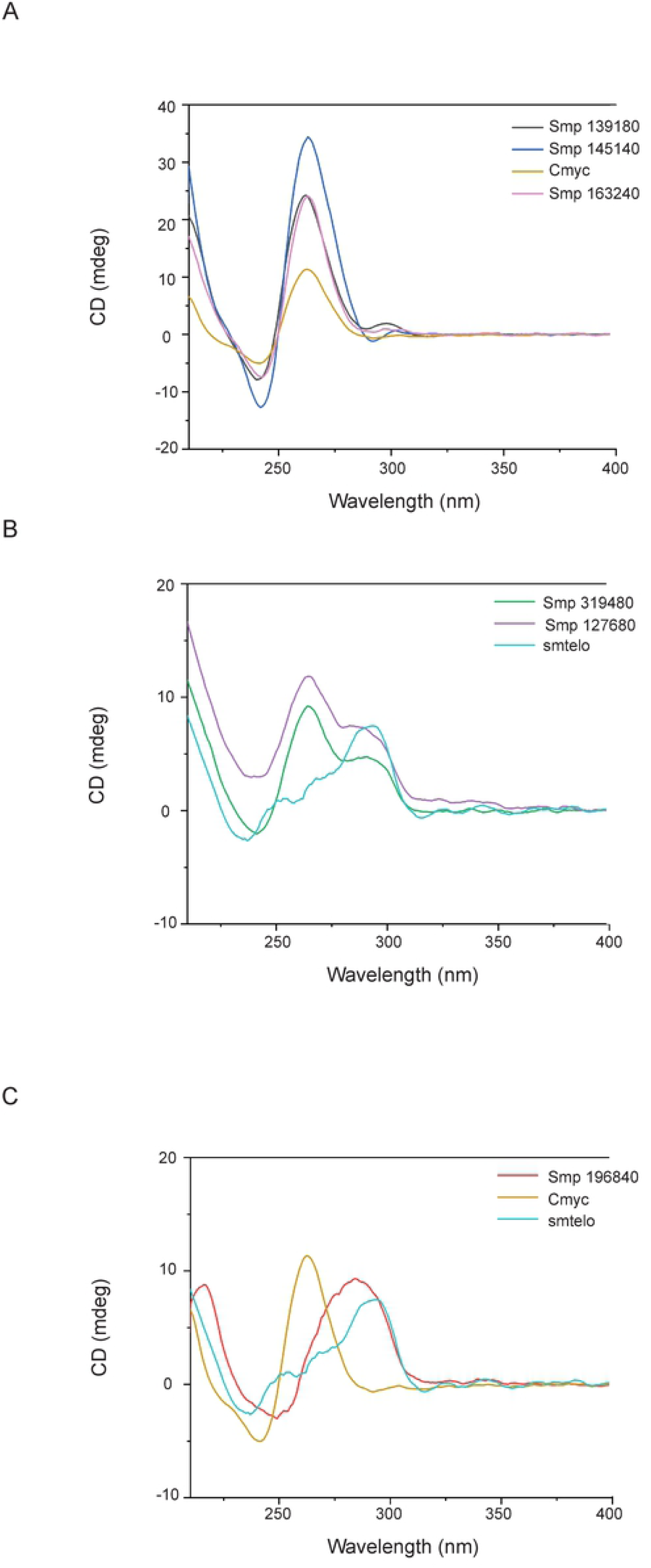
*In vitro* validation of *S. mansoni* oligonucleotides containing PQS by circular dichroism. Oligonucleotides containing predicted PQS within Smps were annealed in the presence of K^+^ before cooling and recording spectra by circular dichroism. A) *smp_139180, smp_145140* and *smp_163240* compared to *c-myc* control, indicating a parallel G4 conformation. B) *smp_319480* and *smp_127680* in comparison to *smTelo* control, with spectra indicative of an antiparallel hybrid quadruplex. C) *smp_196840* with both controls, indicating a spectra readout that does not correlate with either parallel or hybrid quadruplex formation.

Subsequently, melt curves (from 25°C to 95°C) were generated for each of the PQS containing oligonucleotide sequences, in the presence of K^+^, to assess structural stability (S2 Fig.) and the temperature for which half of the initial ellipticity was lost (melting temperature, T_m_) was calculated (S3 Fig). Out of the five tested G4 structures, the oligonucleotide containing the PQS within *smp_145140*, harbouring a parallel G4 conformation, proved to be the most stable as it was not completely melted at 95°C (S2 Fig. B). Interestingly, only *smp_319480’s* representative oligonucleotide underwent a total loss of ellipticity (S2 Fig. E) while, even at 95°C, the other PQSs retained a noticeable secondary structure. Whilst the sequences *smp_139180* (S3 Fig. A), *smp_127680* (S3 Fig. D) and *smp_319480* (S3 Fig. E) share a similar stability (T_m_ ≈ 60 °C), the parallel G4 from *smp_163240* (S3 Fig. C) displayed marked higher stability having a T_m_ of ca. 80 °C.

To assess the role of a cation for proper folding [46], oligonucleotides containing G4 sequences were annealed in the absence of K^+^ and compared to those achieved in presence of 50 mM K^+^ (Fig 5). Interestingly, these CD spectra showed differences in folding profiles and ellipticity in all cases, not corresponding to any type of G4-folding. Overall, in the absence of K^+^, a large decrease in ellipticity as well as a slight blue shift towards lower wavelengths was observed in all spectra.

**Figure 5.**
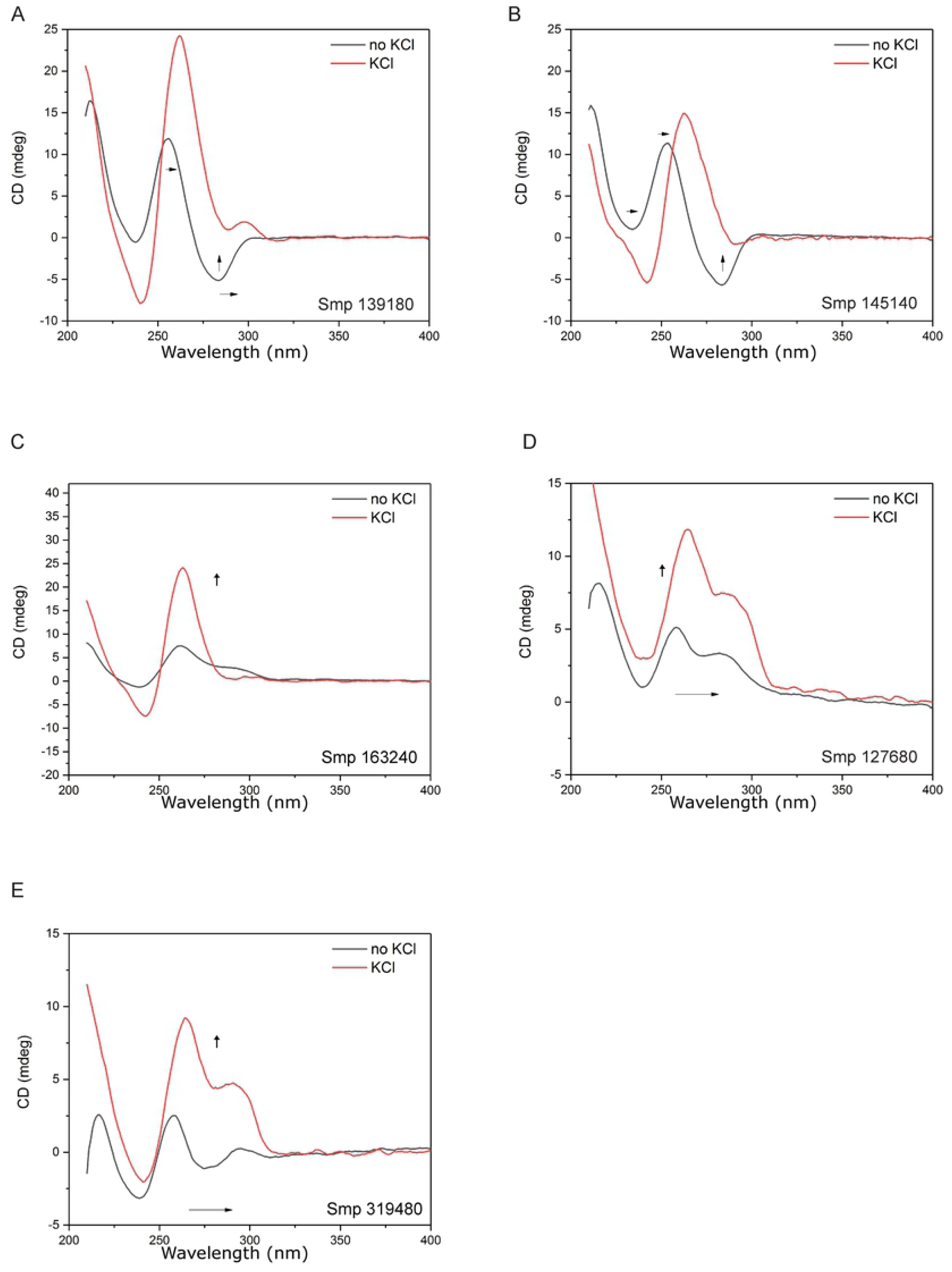
G4 stability of PQS-containing oligonucleotides is dependent on K^+^. PQS containing oligos were annealed in the presence (red line) or absence (black line) of K^+^ ions and spectra recorded for A) *smp_139180* B) *smp_145140* C) *smp_163240* D) *smp_127680* and E) *smp_319480*. Addition of K^+^ ions leads to spectral shifts towards higher wavelengths in all oligonucleotides and increased ellipticity can be observed in the presence of K^+^.

Although CD analyses of 5/6 PQS containing oligonucleotides validated the bioinformatics predictions that G4 structures can be formed within the *S. mansoni* genome and fold into stable conformations, either parallel or hybrid, we wanted to directly identify the presence of G4 epitopes within the parasite itself. Thus, BG4, a single chain variable fragment (scfv) antibody [47], capable of detecting G4 structures in human and *Plasmodium* cells [9,48] was used (Fig. 6).

**Figure 6.**
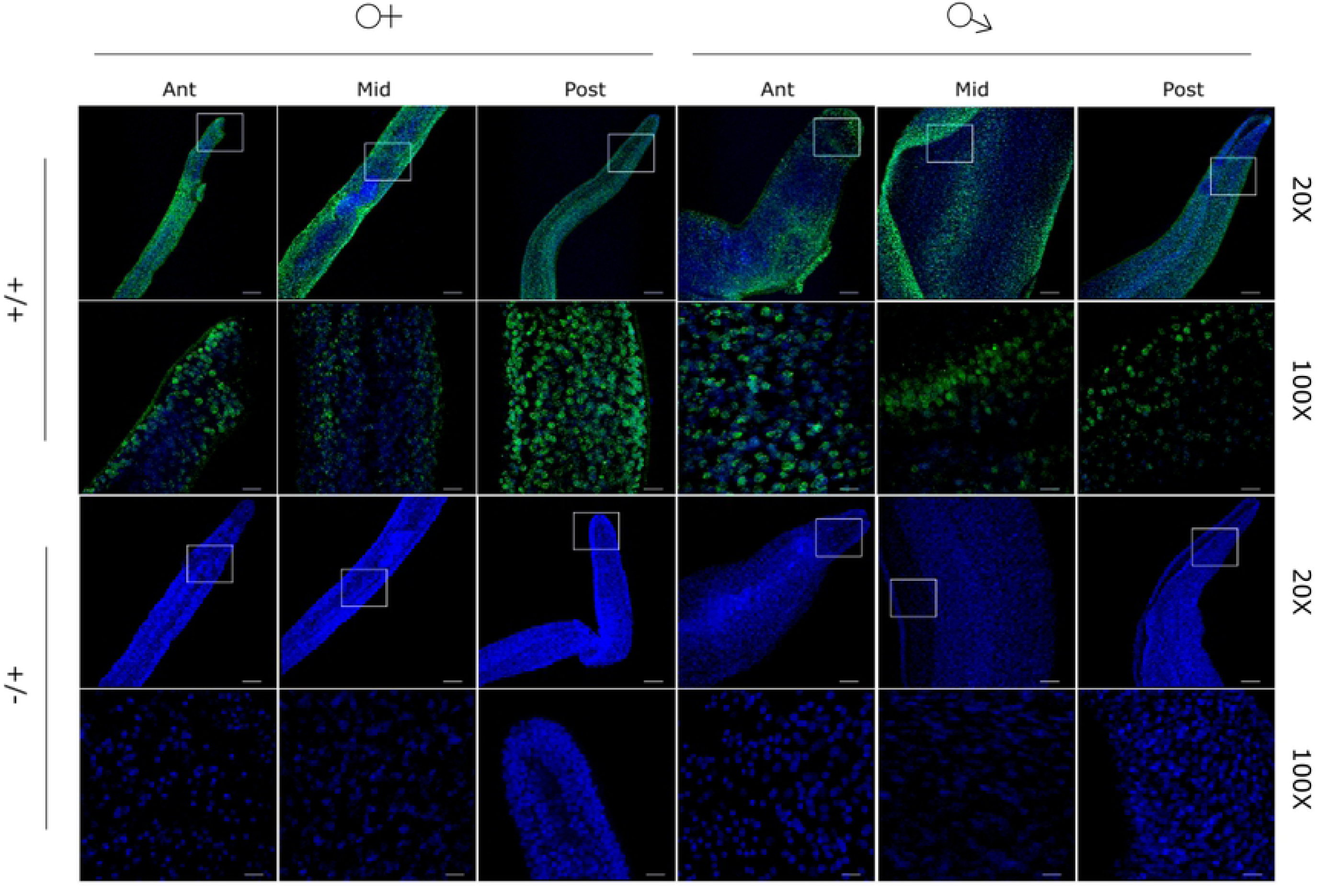
G4 structures are detectable in adult schistosomes. Male and female worms were stained with anti-quadruplex BG4 (green) and counterstained with DAPI (blue). +/+ denotes the use of both primary and secondary antibodies, -/+ denotes the use of secondary antibody only (control). Z stacks of adult anterior (Ant), midsection (Mid) and posterior (Post) regions were taken using a Leica SP8 confocal microscope. Samples were imaged at 20X (scale bar = 50 µm) and then 100X, (scale bar = 10 µm), with box denoting region enlarged. Strong BG4 signal (488 nm) is co-localised with DAPI signal (405 nm) and is observed in both female and male samples.

In male and female worms, BG4 (G4 targeting antibody) signal co-localised with DAPI and was broadly distributed throughout all cell types found within anterior, posterior and midsection regions of both sexes. Loss of BG4 signal was observed when worms were incubated with DNase I prior to processing (Fig. 7A), but not following RNase A incubation (Fig. 7B). As the RNAse A used in these experiments was active (Fig. S4), this data suggested that the vast majority (all) of detectable, adult worm G4 was found in DNA.

**Figure 7.**
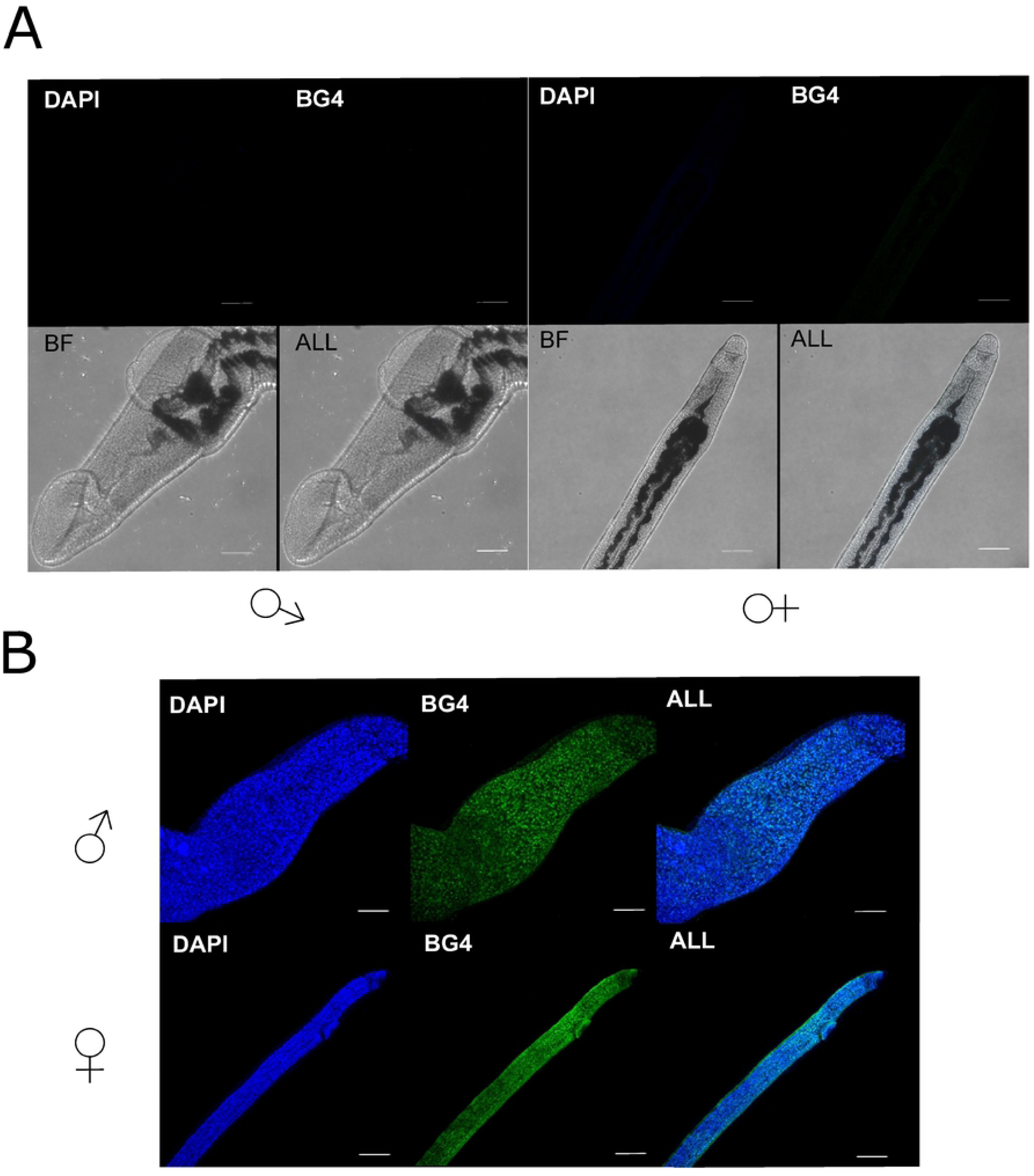
Adult schistosome G4 is predominantly found within DNA, and not RNA, pools. Schistosomes were either treated with A) 120 U DNase I or B) 0.1 mg/ml RNase A for 1 hr at 37°C post permeabilisation and prior to blocking. Anterior sections from males and females were imaged using a Leica SP8 confocal microscope and images collated from Z stacks. Samples were incubated with BG4 (481 nm) and counterstained with DAPI (405 nm) prior to visualisation. Brightfield images were taken for DNase I treated samples due to the absence of any fluorescent signal at either 481 nm (green channel) or blue 405 nm (blue channel). Scale bar denotes 10 µm.

## Discussion

While previous studies have explored the presence and function of G4s in protozoan parasites, similar investigations in multicellular pathogens (i.e. worms) have not been systematically conducted [11,12]. As parasitic helminths remain a significant cause of plant, animal and human disease, new biological insight into how these pathogens regulate complex lifecycles would benefit the development of novel control strategies. Towards this aim, and as an exemplar of a biomedically significant parasitic helminth, we demonstrate that stable G4 structures are present within the *S. mansoni* genome. This finding identifies a previous unexplored area of schistosome biology that may catalyse the development of novel schistosomiasis treatment options and fuel investigations in other parasitic worm species.

Utilising *in silico* approaches involving complementary software tools (QP and G4h), G4 regions are found within both intra- and inter-genic regions of the *S. mansoni* genome (Fig. 2 & 3). Although the abundance of G4s present relative to genome size is low, stringent parameters were enforced to increase detection of true positives; it is, thus, to be expected that more G4 folding sequences exist in the parasite than we have detected here. High-throughput analytical techniques including G4Seq and BG4 CHIP techniques detect a higher incidence of G4 structures compared to computational predictions in a variety of genomes [19,49]. It is, therefore, highly feasible that the same is true for the *S. mansoni* genome. There are also widely reported instances within the literature on sequence based detection of non-classical G4 structures (e.g. bulge quadruplexes or loop errors/mutations) that are not detectable by QP and are limited in detection by G4h [50,51]. Therefore, many non-classical G4 would be missed in our analyses and this will also contribute to a conservative initial estimate.

The high number of PQS located in intergenic regions compared to intragenic regions is largely due to the tandem telomeric repeat widely confirmed to adopt G4 folding [52–54]. A search for the ‘TTAGGG’ repeat (and its reverse complement to account for the reverse strand) motif within PQS found that a large majority of intergenic PQS to be telomeric, a trend found across all chromosomes but in particular chromosome 5 (S1 Table). This may simply be due to quality of sequence coverage of chromosome 5 in the current genome assembly compared to other chromosomes. Telomeric PQS comprising the majority of detected intergenic PQS has been observed in *P. falciparum* and indeed many eukaryotes [12,16,19]. While non telomeric intergenic PQS have been observed across many species, research into their function is mostly limited to intragenic counterparts [9,55,56]. Non-telomeric intergenic PQS have been hypothesised to block transcription in *Arabidopsis* due to their proximity to transcription start sites, but this has not been explored greatly in other systems [56].

Although the range of intragenic PQS across chromosomes is fairly limited (7-23 PQS), chromosome 5 appears greatly enriched for PQS compared to all other chromosomes following normalisation (Fig 3). Chromosome 5 does not differ significantly in GC content compared to other chromosomes (Fig 2B) and indicates that PQS are not present due to an increase in GC content in the chromosome. The reason for increased numbers of PQS in this chromosome is currently unclear and requires further exploration.

The low absolute number of PQS detected within exons may, in part, be due to the phenomenon of large introns within the genome [2]. Indeed, the average length of an *S. mansoni* intron is 1,692 bp (with some up to 33.8 kb reported) compared to an average exon length of 217 bp and this likely accounts for a higher number of absolute intronic PQS compared to other intragenic motifs [2]. Upon normalisation, the intragenic PQS rate is scarce, but it is important to note that *S. mansoni* has a relatively large genome (397.2 Mb) with long stretches of noncoding areas and this trait is also observed in other AT rich genomes such as *Plasmodium* [3,16]. Despite a high absolute number of intronic PQS, the greatest enrichment of PQS (when normalised to chromosomal feature) was detected in the 3’ regions, and then 5’ regions. Mammalian genomes (mouse and human) are also enriched for PQS within the 5’ region and especially in promoters [19], but in contrast do not feature the strong 3’ enrichment seen here in *S. mansoni*. Over-enrichment of PQS in 3’ UTRs has also been observed in the *Arabidopsis* genome, although the functionality of such motifs was not explored further [19]. G4s within mRNA 3’ UTRs are associated with cis-regulatory control mechanisms including translational repression and regulation of miRNA binding, with G4s masking miRNA binding sites and leading to shorter transcripts through modulation of polyadenylation efficiency [57,58]. The *S. mansoni* genome encodes miRNAs with research suggesting an important role in parasite developmental biology, sexual maturation and host cell modulation [59–61]. G4s in 3’ UTRs may thus, potentially be associated with miRNA mediated gene expression regulation within the parasite, highlighting a future endeavour to be explored in further contextualising the functional role of *Schistosoma* G4.

GO term analysis revealed a significant enrichment of PQS within genes associated with the *wnt* signalling pathway (confirmed by CD for *smp_145140*-*wnt* and *smp_139180*- *frizzled*; Fig. 4), a crucial developmental system necessary for parasite development [62–64]. In the Platyhelminthes, *wnt* is associated with proper axial development and maintenance of anterior-posterior polarity [4,64] and so targeting G4 present within *wnt*-associated genes may provide a potential avenue in disrupting parasite development. *S. mansoni* contains a *wnt* gene orthologous to planarian *wnt2* that is expressed in a spatial gradient along the worm posterior, suggesting linkage with axial development [4]. Within *S. japonicum*, both *wnt* (*Sjwnt4* and *Sjwnt5*) and *frizzled (Sjfz7*) genes have been identified and characterised across multiple lifecycle stages of the parasite [65–67]. *Sjwnt5* was additionally linked with reproductive organ development and is found highly expressed in testes, ovaries and vitellarium [68]. Interestingly, G4s have been identified in genes encoding components of the *wnt* pathway in other organisms. Specifically, G4s are found in the promoter region of human *wnt1* and this was found to be sensitive to G4 stabilising compounds [69]. Furthermore, G4s have also been described in both human and zebrafish *fzd5* where they have been associated with transcriptional control of the gene [70,71]. Collectively, these findings suggest that G4 enrichment in genes encoding *wnt* pathway components is conserved across genomes and may have functional implications.

CD spectra demonstrate that PQS within *S. mansoni* protein coding genes can fold unambiguously into higher order G4 structures when exposed to quadruplex favouring conditions (Fig. 4). PQS-containing oligonucleotides representing *smp_139180, smp_163240* and *smp_145140* folded into structures with spectra comparable to the *H. sapiens c-myc* control, a known parallel G4. In contrast, PQS-containing oligonucleotides representing *smp_319480* and *smp_127680* displayed CD spectra that mirrored that of hybrid G4s found in *smTelo* as well as elsewhere [72–74], and showed a comparable stability. With respect to the parallel folded G4s, the high T_m_ observed for the oligonucleotide *smp_163240* and the impossibility to extrapolate a T_m_ for *smp_145140*, indicate thermodynamic stability associated with single nucleotide loop motifs (Fig. S3), which have been characterised in *H. sapiens vegf* and *c-kit*, as well as the *c-myc* control used in our study [75,76]. Indeed, the only oligonucleotide representative to adopt a parallel folded G4 structure and to not contain a single nucleotide loop was derived from *smp_139180*; this oligonucleotide also demonstrated the lowest T_m_ among the three sequences.

Of note, the only PQS-containing oligonucleotide that did not resolve into a G4 structure was derived from *smp_196840*. Specifically, this oligonucleotide displayed CD spectra more commonly associated with B form DNA structures [77]. A possible explanation for the lack of classical G4-folding for this DNA sequence, when compared to the others, could be due to the length of the sequence; *smp_196840*’s oligonucleotide was the second longest (33 nt) and contained the longest loop length between G quartets (9 nt). Combined, these two factors may contribute to a greater instability of the oligo and a reduced likelihood of G4 formation, as shorter loop lengths are often associated with a more stable tertiary structure [78,79]. This phenomenon is not unusual, with other reported instances of computationally predicted PQS not folding *in vitro* [78].

The CD data also confirm that G4 folding is K^+^ dependent (Fig. 5). Cations are crucial for G4 formation and stability, contributing to G4 tetrad stabilisation by electrostatic binding to the O6 of the guanines and interacting with the negative charges of the phosphate backbone [80]. Although there is some higher order folding detectable in the *S. mansoni* G4 sequence in absence of monovalent cations (Fig. 6), these are likely not G4-type structures, as indicated by the spectra and ellipticity. Therefore, K^+^ is needed for G4 folding within *S. mansoni* derived PQS.

The immunofluorescent detection of G4 within adult worm nuclei (DAPI^+^) clearly validates the *in silico* predictions/CD spectra confirmation of PQS within this parasitic worm (Fig. 6). While the BG4 antibody can recognise G4 epitopes in both RNA and DNA [81], the loss of signal when adult worms were treated with DNase I, but not RNaseA, strongly suggests that most G4s are found in the DNA pool (Fig. 7). Within the nucleus, the location of the DNA-associated G4 cannot be specifically resolved by our current LSCM analyses. However, the transient nature of G4s and the abundance required for visualisation by this technique, it is likely that the signal observed here is derived from telomeric G4s given the large number detected in computational analyses (S1 Table). As this indicates that many PQS are present within the terminal ends of chromosomes (derived from the schistosomal telomeric tandem repeat TTAGGG) and this repeat can form a stable G4 structure (Fig 4) similar to *htelo* [82– 85], this reasonably accounts for the signal seen. However, detection of G4 in metaphase chromosomes could provide further information relevant to spatial PQS distribution in *S. mansoni*. Equally, exploration of BG4 signal localisation at different cell cycle phases could also further contextualise the profile of BG4 signal.

In summary, G4s were identified in the parasitic platyhelminth *S. mansoni* for the first time. Computational analysis suggests that G4s are distributed across all chromosomes with *in vitro* validation that putative G4-containing sequences can fold into stable PQS as visualised by CD spectroscopy. Confocal microscopy of adult schistosomes demonstrates the *in vivo* presence of G4s in fixed parasites. This lends credence to the presence of G4 as functional operators within the parasite and opens up previously unknown avenues for exploration in parasite biology, development and regulation.

## Acknowledgements

We acknowledge Ms Julie Hirst and all members of the Hoffmann laboratory for assistance in *Schistosoma* life cycle maintenance. We thank Dylan Phillips and Alan Cookson for advice on fluorescence imaging. Some *B. glabrata* snails used in this study were provided by the NIAID Schistosomiasis Resource Centre of the Biomedical Research Institute (Rockville, MD, USA) through NIH-NIAID Contract HHSN272201000005I for distribution through BEI Resources. This work was supported by Aberystwyth University, the Wellcome Trust (107475/Z/15/Z) and The Joy Welch Educational Charitable Trust.

## Author Contribution

Conceptualization: HC, HW, MS, AC, and KFH; Formal Analysis: HC, VL, RB, MS, DB; Funding Acquisition: HW, KFH; Investigation: HC, RB, NS; Project Administration: KFH, AC; Resources: KFH, AC; Software: MS, VL Supervision: KFH, HW, MS; Visualisation: HC, RB; Writing – Original Draft Preparation: HC; Writing – Review and Editing: HC, HW, KFH

**S1 Figure. Analysis of GO terms in PQS containing Smps**. A) GOAtools analysis was performed on GO terms from Smps that had computationally predicted PQS. Output was plotted and *p* < 0.05 indicated by dashed line. GO terms that had significant enrichment included GO:0016055 – Wnt signalling pathway, GO:1905114 – cell surface receptor signalling pathway involved in cell-cell signalling and GO:0005201 – extracellular matrix structural constituent. B) A ReVigo tree analysis of the same GO terms. *Wnt* signalling is the most enriched molecular function related GO term.

**S2 Figure. Structure of PQS containing oligonucleotides are differentially affected by temperature**. Oligos were adjusted to a 6 µM working solution in 60 mM TrisKCl (10 mM TrisHCl, 50 mM KCl; pH 7.4) buffer. CD spectra were recorded at 5°C intervals between 25°C and 95°C; data was used to calculate T_m_s. Parallel forming oligonucleotide sequences A) *smp_139180*, B) *smp_145140* and C) *smp_163240* showed stable structures that lost some ellipticity but not completely. Hybrid G4 folding oligonucleotide sequences D) *smp_127689* followed the same trend as above whereas E) *smp_319480* totally lost its ellipticity as temperature increased, indicating a complete unfolding of the G4.

**S3 Figure. Melt curves of thermodynamic stability analysis**. Spectra was recorded by CD at 5 °C intervals between 95 and 25 °C. Normalised CD was plotted, and T_m_ determined for each oligo sequence. Three repeats (average reading shown) were performed for each temperature point and regression analysis was performed to fit curves.

**S4. Figure RNaseA is enzymatically active**. A sample of 1 µg each of total murine RNA and *S. mansoni* RNA were incubated for 1 hr in 0.1 mg/ml RNase (37 °C) and electrophoresed on an agarose gel with a 1kB ladder alongside RNA only controls (incubated under the same conditions in the absence of RNaseA). Complete degradation of host/parasite RNA was only observed in the presence of RNaseA (R+).

